# A Decoy Library Uncovers U-box E3 Ubiquitin Ligases that Regulate Flowering Time in Arabidopsis

**DOI:** 10.1101/2020.03.02.973149

**Authors:** Ann M. Feke, Jing Hong, Wei Liu, Joshua M. Gendron

**Affiliations:** Department of Molecular, Cellular, and Developmental Biology, Yale University, New Haven, CT 06511, USA; School of Food Science and Engineering, South China University of Technology, Guangzhou, China

## Abstract

Targeted degradation of proteins is mediated by E3 ubiquitin ligases and is important for the execution of many biological processes. Previously, we created and employed a large library of E3 ubiquitin ligase decoys to identify regulators of the circadian clock (Feke et al., 2019). In tandem with the screen for circadian regulators, we performed a flowering time screen using our U-box-type E3 ubiquitin ligase decoy transgenic library. We identified five U-box decoy transgenic populations that have defects in flowering time or the floral development program. We used additional genetic and biochemical studies to validate *PLANT U-BOX 14* (*PUB14*), *MOS4-ASSOCIATED COMPLEX 3A (MAC3A),* and *MAC3B* as *bona fide* regulators of flowering time. This work reinforces the utility of the decoy library in identifying regulators of important developmental transitions in plants and expands the scope of the technique beyond our previous studies.

## INTRODUCTION

Flowering is the first committed step in the plant reproductive process, leading to the production of reproductive organs and eventually offspring. Plants use highly complex gene networks that integrate a wide array of internal and external signals to regulate flowering. In the model plant *Arabidopsis thaliana*, six pathways have been identified to control flowering time. Four of these six pathways regulate the production of the florigen, the protein FT, while the remaining two bypass FT to promote flowering in a more direct manner (Srikanth and Schmid, 2011). In addition, abiotic and biotic stress can modulate flowering time by altering the function of one or multiple flowering pathways (Park et al., 2016; Takeno, 2016).

In Arabidopsis, the transition to flowering is an irreversible decision the plant makes in response to external and internal signals. In order for the response to occur at the appropriate time, plants need to promote the activity of floral activators and repress the activity of floral repressors. One way that plants accomplish this is by leveraging the ubiquitin proteasome system to accurately degrade floral regulator proteins (Hu et al., 2014; Imaizumi et al., 2005; McGinnis et al., 2003; Nelson et al., 2000; Sawa et al., 2007). E3 ubiquitin ligases provide substrate specificity for the ubiquitin proteasome system and mediate ubiquitylation of target proteins (Vierstra, 2009). E3 ubiquitin ligases play central roles in the regulation of the photoperiodic, vernalization, and GA flowering time pathways (Hu et al., 2014; Imaizumi et al., 2005; Jang et al., 2008; Lazaro et al., 2012; McGinnis et al., 2003; Nelson et al., 2000; Sawa et al., 2007) demonstrating their important functions in this critical developmental decision.

The study of E3 ubiquitin ligases in plants can be difficult due to the numerous genome duplications that have resulted in widespread functional redundancy. To overcome this we previously created a library of transgenic plants expressing E3 ubiquitin ligase decoys (Feke et al., 2019). We tested the effects of the E3 ubiquitin ligase decoys on the circadian clock and found dozens of new potential clock regulators. We then performed follow-up studies on two redundant U-box genes, MOS4-ASSOCIATED COMPLEX 3A AND MOS4-ASSOCIATED COMPLEX 3B (MAC3A and MAC3B), two redundant U-box genes that control splicing of a core circadian clock transcription factor. We proposed that this library could be used to study any biological process of interest in Arabidopsis.

The UPS plays known roles in the control of flowering time (Hu et al., 2014; Imaizumi et al., 2005; Jang et al., 2008; Lazaro et al., 2012; McGinnis et al., 2003; Park et al., 2007; Sawa et al., 2007; Song et al., 2012; Sun, 2011). Despite this, it is possible that E3 ubiquitin ligases that regulate flowering have not been identified due to the aforementioned gene redundancy (Navarro-Quezada et al., 2013; Risseeuw et al., 2003; Yee and Goring, 2009). Here, we employ the decoy library to identify U-box-type E3 ubiquitin ligases that control flowering time and reproductive development. We focus on four metrics of reproductive development: the number of leaves at 1 cm bolting, the age of the plant in days when 1 cm bolting occurs, the first occurrence of anthesis, and the rate of stem elongation. Using these metrics, we uncover six U-box proteins which regulate 1 cm bolting, six U-box proteins which regulate leaf number, four U-box proteins that control stem elongation, and one U-box protein that controls anthesis.

We perform focused genetic studies on three U-box genes, *PLANT U-BOX 14* (*PUB14*), *MAC3A,* and *MAC3B.* We confirm their roles in flowering time regulation by observing delayed flowering phenotypes in three T-DNA insertion mutants, *pub14-1* (SALK_118095C), *mac3a,* and *mac3b* mutants (Monaghan et al., 2009). We also perform immunoprecipitation mass spectrometry with the PUB14 decoy, similar to what we did previously for MAC3B (Feke et al., 2019), and find a list of proteins involved in the regulation of flowering time. These findings build on our previous manuscript by showing that the decoy library can be effective for identifying E3 ubiquitin ligases that participate in plant developmental processes outside of the circadian clock, and illustrate the strength of the decoy technique to quickly identify novel E3 ubiquitin ligases in diverse biological processes.

## RESULTS

### The Role of U-box Decoys in Flowering Time

Protein degradation through the ubiquitin proteasome system plays an essential role in flowering time pathways (Hu et al., 2014; Imaizumi et al., 2005, 2003; Jang et al., 2008; Lazaro et al., 2012; McGinnis et al., 2003; Park et al., 2007; Sun, 2011). However, the extent to which the ubiquitin proteasome system regulates flowering is not fully known. In order to identify E3 ubiquitin ligases that regulate flowering time, we screened the U-box decoy library. This is a subset of the larger decoy library described in our previous manuscript (Feke et al., 2019).

Parental control and T1 transgenic seedlings expressing the decoys were transferred to soil and grown under long day (16 hours light, 8 hours dark) conditions. By analyzing a population of T1 transgenics, we avoid the problems that may arise from following a single insertion that may not be representative of the entire population.

In order to monitor the initiation of flowering, we measure the number of leaves at 1 cm bolting. This is a common flowering time measurement that informs on the developmental stage of the plant at the vegetative to reproductive phase transition. We also measure the age of the plant in days when bolting occurs, as indicated by a 1 cm long inflorescence. This allows us to determine how much time the plant spends in the vegetative stage.

In addition to floral initiation, we also measure two metrics of reproductive development. We measure the first occurrence of anthesis, or the opening of the floral bud. While Arabidopsis self-pollinates prior to anthesis, anthesis is required for fertility in plants that are pollinated externally (Khanduri, 2011; Rivero et al., 2014). As anthesis is dependent on the initiation of flowering, we calculated the anthesis delay, or number of days after 1 cm bolting that anthesis occurs, and used this value for our analyses. We also measure stem elongation by recording the age of the plant in days when the inflorescence is 10 cm long. Stem elongation may be a measure of fertility as it is correlated with the appearance of internodes (Carvalho et al., 2002). Similar to anthesis, stem elongation depends on the initiation of flowering. Thus, we calculate the stem elongation time, or the difference between the age of the plant at 10 cm stem length and the age at the plant at 1 cm stem length. By measuring all four metrics, we are able to categorize any candidate floral regulator by which aspects of floral development are impacted.

Flowering time and reproductive development can differ between experiments due to uncharacterized variations in growth conditions. In order to compare across the entire decoy library, we calculated the flowering time difference for each individual decoy transgenic. This value was calculated by determining the average leaf number for the control population in each experiment, then subtracting this value from the leaf number for each individual decoy transgenic (Figure 1). We generate the 1 cm bolting time difference (Figure 2), anthesis delay difference (Figure 3), and stem elongation period difference (Figure 4) in this same manner. In order to see the variation within experiments, the individual control plants were normalized against the other control plants in the same experiment, as described for the decoy plants above (Figure 1 – Figure Supplement 1, Figure 2 – Figure Supplement 1, Figure 3 – Figure Supplement 1, and Figure 4 - Figure Supplement 1). We perform a Welch’s t-test with a Bonferroni-corrected α of 1.25×10^-3^ on these difference values. In this way, we are able to confidently assess whether a decoy population is different from the control in any of our metrics.

**Figure 1.**
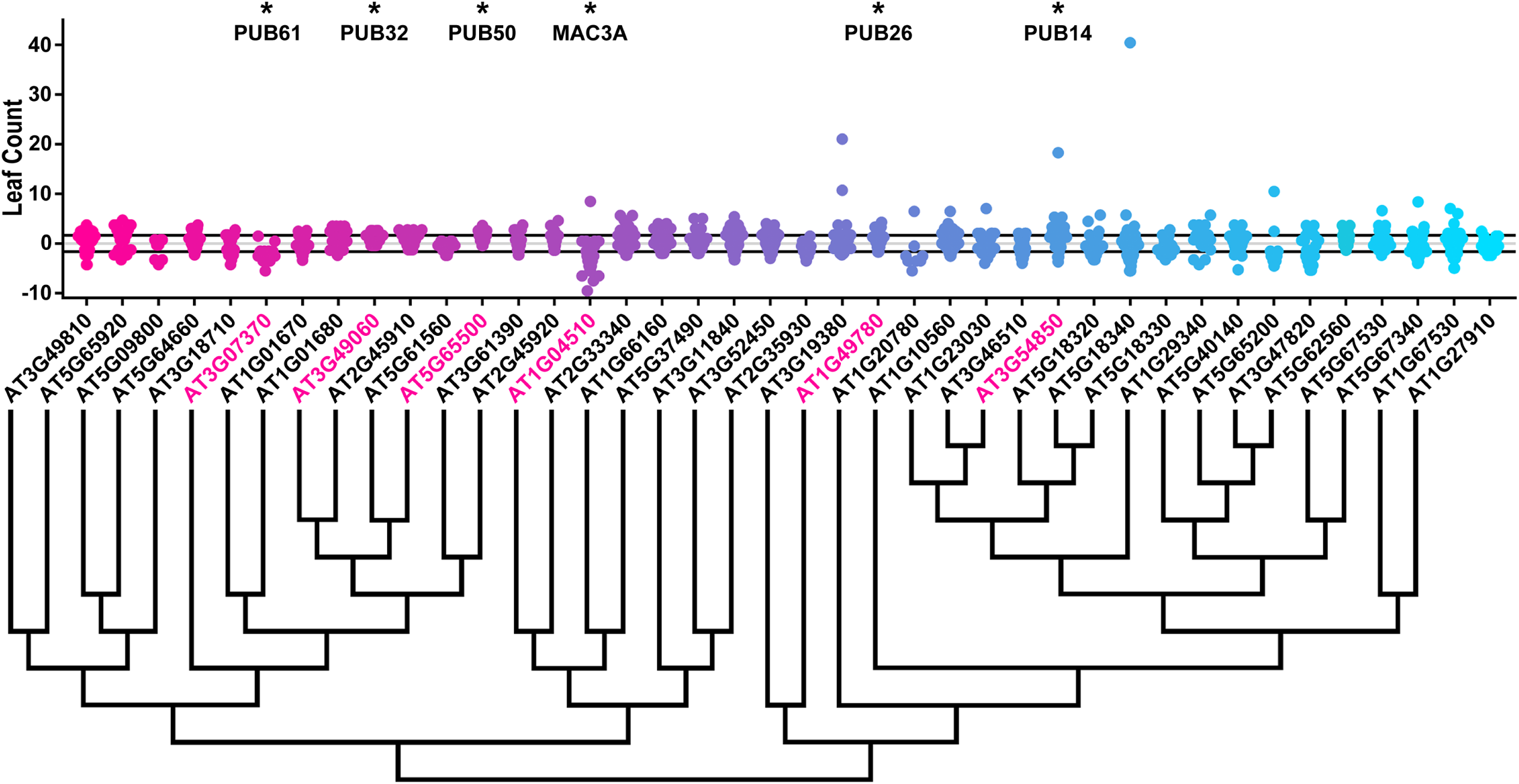
Leaf Count Distributions of U-box Decoy Plants. Values presented are the difference between the leaf count 1 cm inflorescence of the individual decoy plant and the average leaf count at 1 cm inflorescence of the parental control in the accompanying experiment. The grey line is at the average control value and the black lines are at +/- the standard deviation of the control plants. Genes are ordered by closest protein homology using Phylogeny.Fr, (Dereeper et al., 2008), and a tree showing that homology is displayed beneath the graph. * and pink gene names = The entire population differs from wildtype with a Bonferroni-corrected *p* <1.25×10^-3^.

**Figure 2.**
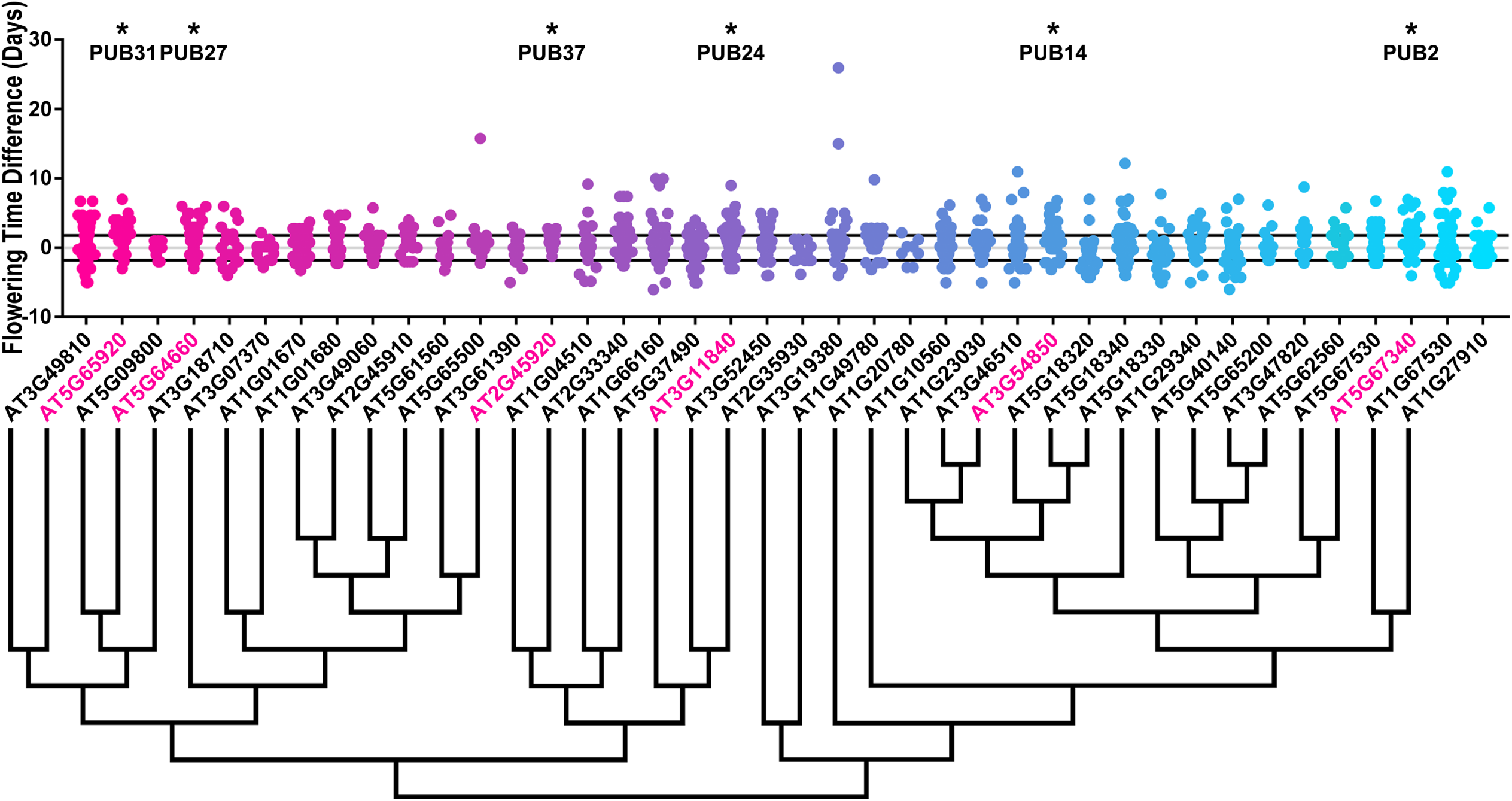
1 cm Bolting Age Distributions of U-box Decoy Plants. Values presented are the difference between the age at 1 cm inflorescence of the individual decoy plant and the average age at 1 cm inflorescence of the parental control in the accompanying experiment. The grey line is at the average control value and the black lines are at +/- the standard deviation of the control plants. Genes are ordered by closest protein homology using Phylogeny.Fr, (Dereeper et al., 2008), and a tree showing that homology is displayed beneath the graph. * and pink gene names = The entire population differs from wildtype with a Bonferroni-corrected *p* <1.25×10^-3^.

**Figure 3.**
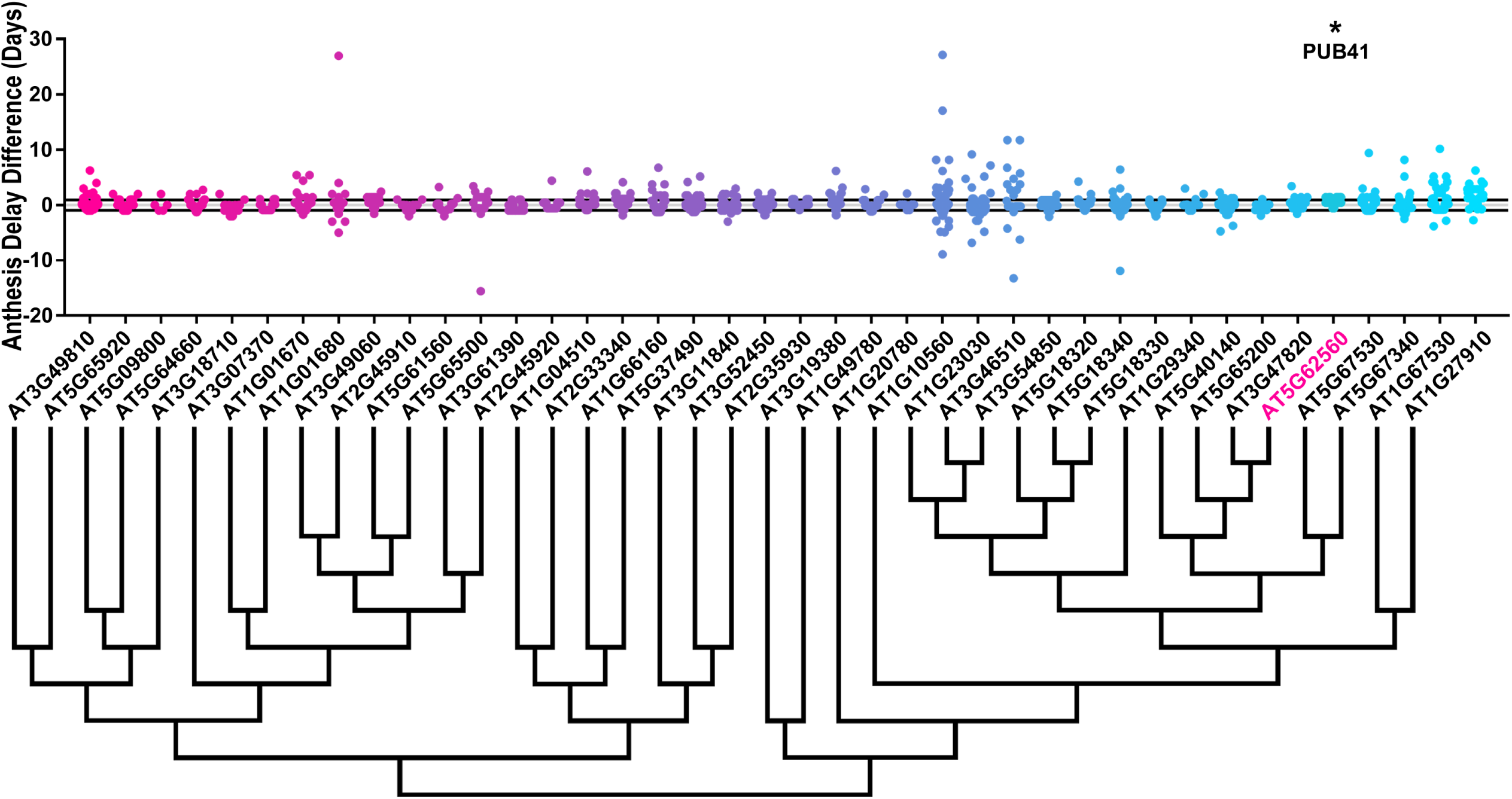
Anthesis Delay Distributions of U-box Decoy Plants. Values presented are the difference between the anthesis delay of the individual decoy plant and the average anthesis of the parental control in the accompanying experiment. Anthesis delay is defined as number of days between the inflorescence height reaching 1 cm and the first flower bud opening. The grey line is at the average control value and the black lines are at +/- the standard deviation of the control plants. Genes are ordered by closest protein homology using Phylogeny.Fr, (Dereeper et al., 2008), and a tree showing that homology is displayed beneath the graph. *and pink gene names = The entire population differs from wildtype with a Bonferroni-corrected *p* <1.25×10^-3^.

**Figure 4.**
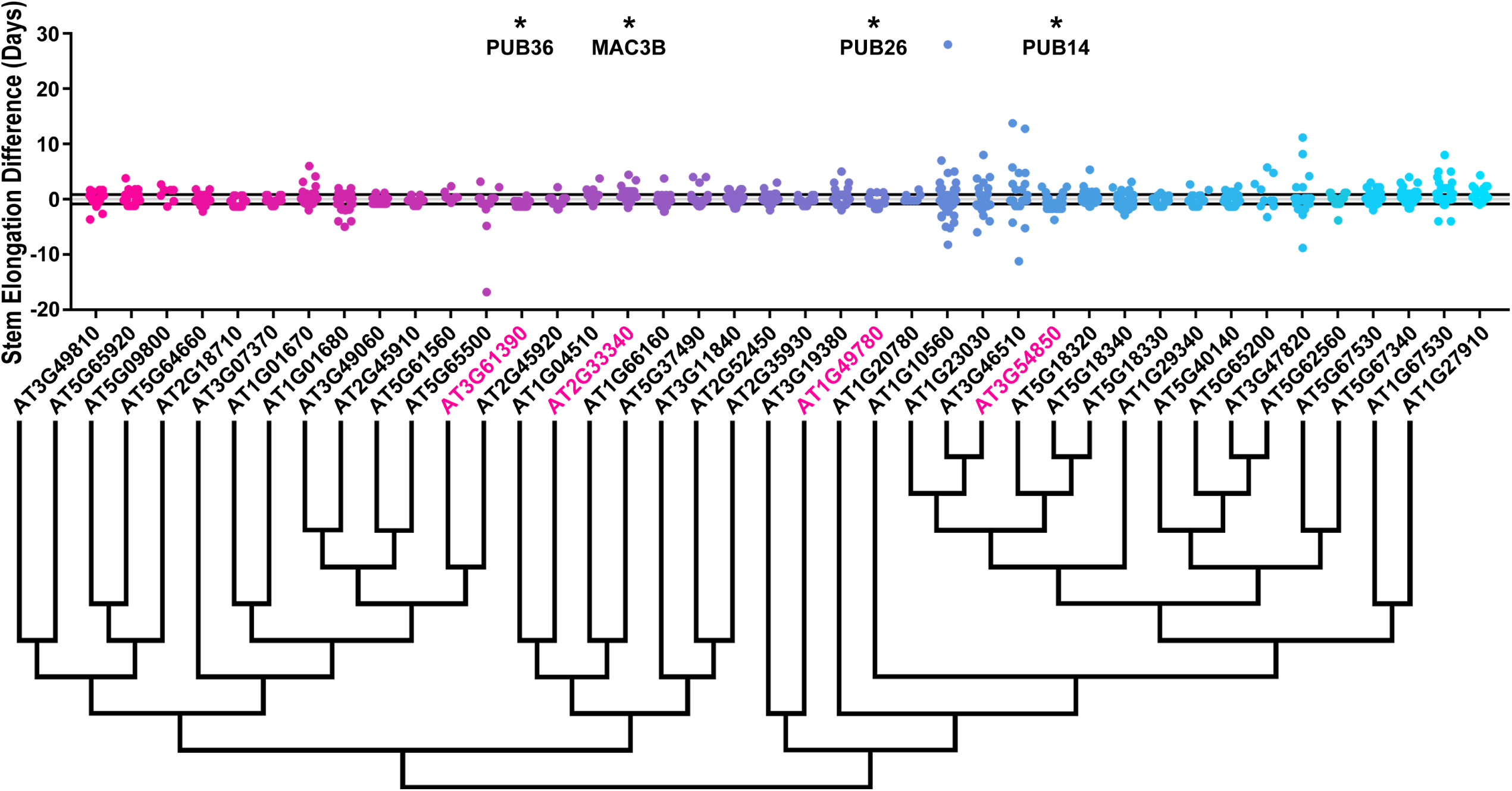
Stem Elongation Time Distributions of U-box Decoy Plants. Values presented are the difference between the stem elongation period of the individual decoy plant and the average stem elongation period of the parental control in the accompanying experiment. The stem elongation period is defined as the number of days between the inflorescence height reaching 1 cm and the inflorescence height reaching 10 cm. The grey line is at the average control value and the black lines are at +/- the standard deviation of the control plants. Genes are ordered by closest protein homology using Phylogeny.Fr, (Dereeper et al., 2008), and a tree showing that homology is displayed beneath the graph. * and pink gene names = The entire population differs from wildtype with a Bonferroni-corrected *p* <1.25×10^-3^.

Of the 40 decoy populations assayed, six populations demonstrated a statistically altered age at 1 cm bolting, six had altered leaf number, one had altered anthesis, and four had altered stem elongation time. Most effects on flowering time were minor. We define “minor” as less than two leaves different than wild type at 1 cm bolting or less than two days different than wildtype in 1 cm bolting, anthesis, or stem elongation time, and “major” as more than two leaves or days different from wildtype, respectively. To narrow candidates for detailed genetic follow-up studies we focused on decoy populations that had any effect on multiple flowering criteria, or had a major effect on one criterion.

There were three decoy populations that had a major effect on one flowering criterion. Expressing the *PUB31* decoy had a large effect on 1 cm bolting, delaying it by an average of 2.4 days. Expressing *MAC3A* and *PUB61* decoys caused altered leaf number at 1 cm bolting with 2.9 and 2.2 fewer leaves, respectively. The magnitude of the phenotype observed in these populations makes them high-priority candidate flowering time regulators. *MAC3A* had the greatest magnitude change in leaf number at 1 cm bolting, was previously noted to have a flowering time defect (Monaghan et al., 2009), and was part of our focused circadian clock genetic studies previously (Feke et al., 2019) making it a major candidate for focused genetic studies with regards to its role in flowering time.

In order to identify additional major candidate flowering time regulators, we determined which decoy populations caused defects in multiple flowering time parameters (Figure 5). Expressing the *PUB26* decoy shortens the stem elongation period by 0.75 days, and results in 1.4 more leaves at flowering time. This population may also have delayed 1 cm bolting by 1.7 days, but this difference did not reach our strict statistical cutoff (*p* = 0.036). *PUB14* also affects multiple flowering parameters. It shortens the stem elongation period (0.56 days shorter), flowers with more leaves (1.3 leaves), and also delays 1 cm bolting (1.8 days), with all three parameters reaching statistical significance. Many classic flowering time regulators affect both leaf number at 1 cm bolting and days to 1 cm bolting (Nelson et al., 2000; Page et al., 1999; Wang et al., 2011), making PUB14 a strong candidate for follow-up studies.

**Figure 5.**
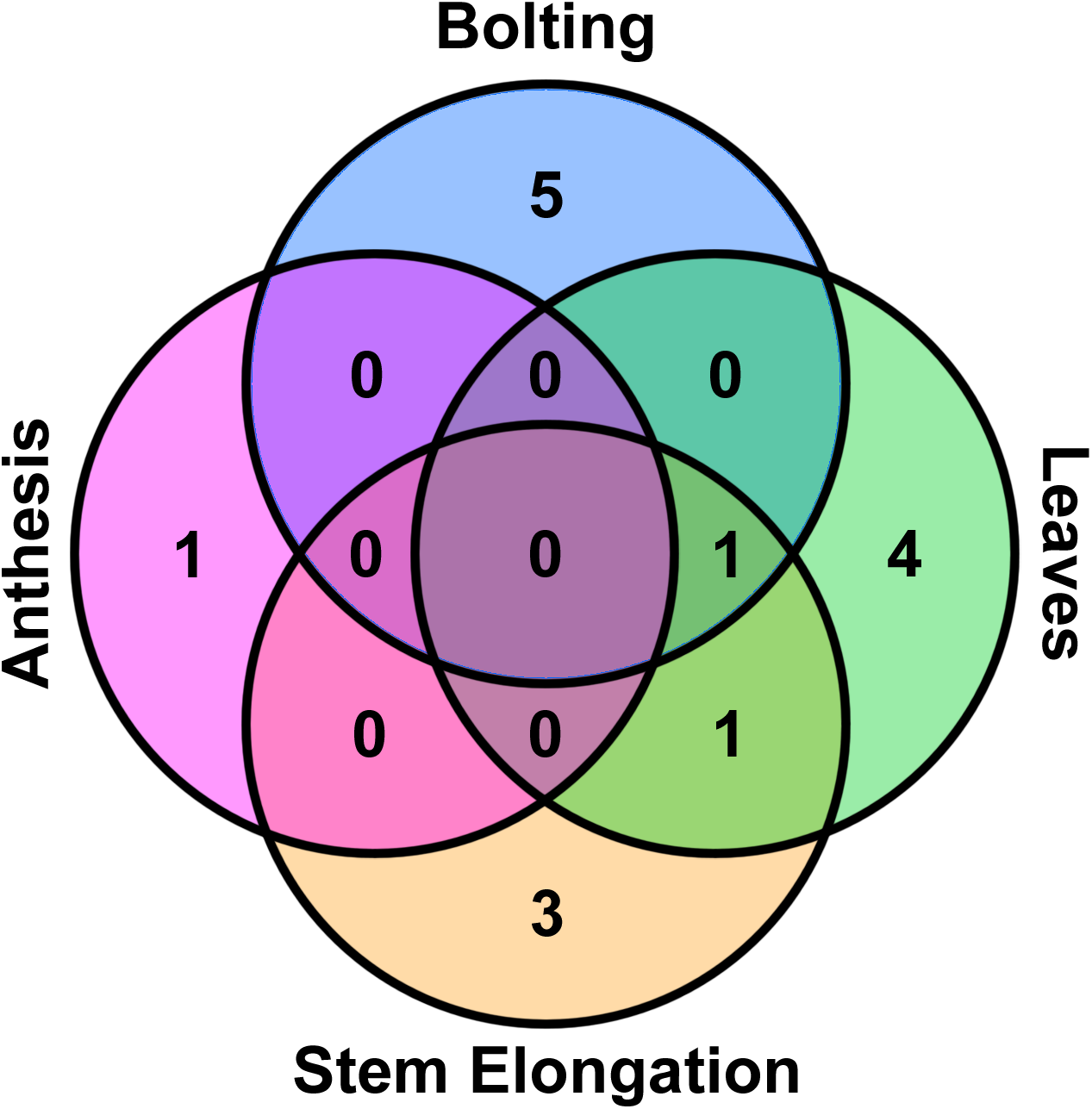
Overlap Between Candidate Flowering Time Regulators for each Metric. The statistically significant regulators from Figures 1-4 were categorized based on which metrics were affected.

### *PUB14* Regulates Flowering Time

PUB14 was the only candidate flowering time regulator identified in our screen that impacted both leaf number and 1 cm bolting age. However, the function of *PUB14* in flowering time regulation is unknown. In order to understand the function and regulation of *PUB14*, we mined publically available expression data and the literature. PUB14 has the U-box domain centrally located and possesses five ARMADILLO (ARM) repeats. It has no known genetic function, although it was used as a “prototypical” *PUB* gene in a structural study on U-box function (Andersen et al., 2004). *PUB14* is closely related to *PUB13* (E-value of 0), which has been implicated in the control of flowering time, immunity, cell death, and hormone responses (Kong et al., 2015; Li et al., 2012b, 2012a; Liao et al., 2017; Zhou et al., 2018, 2015). Mutants of *PUB13* have accelerated flowering time in long day conditions (Li et al., 2012b, 2012a; Zhou et al., 2015), in contrast to the delayed flowering we observed with the *PUB14* decoys. Although this data suggests that *PUB14* and *PUB13* affect flowering time differently, the identification of a close homolog of a characterized flowering time regulator provides strength to the hypothesis that *PUB14* could also regulate flowering time.

Genes that regulate flowering time are often regulated by the circadian clock or diel light cycles, and their expression may differ between inductive (long day) and non-inductive (short day) conditions. Thus, we attempted to determine whether *PUB14* is regulated by the circadian clock or various daily light cycles. In order to assay for rhythmicity, we queried publically available microarray data and determine the correlation value, a measure of the similarity between the expression data and the hypothesized cycling pattern (Mockler et al., 2007). If this correlation value is greater than the standard correlation cutoff of 0.8, then it is considered rhythmic. While *PUB14* expression does not cycle under circadian, LD (12 hour light/12 hour dark) or floral inductive long day (16 hour light/8 hour dark) conditions, it does cycle under non-inductive short day (8 hour light/16 hour dark) conditions, peaking in the evening (19 hours after dawn) (Mockler et al., 2007). Furthermore, many flowering time genes are regulated by stress, temperature, or hormones. For this reason, we mined expression data using the eFP browser for treatments that effect *PUB14* expression (Winter et al., 2007). While *PUB14* expression is unaffected by most treatments, it is upregulated when leaves are exposed to *Pseudomonas syringae* (Winter et al., 2007).

*PUB14* is closely related to a gene that regulates flowering time and expressing the decoy causes delayed flowering. We isolated an Arabidopsis mutant with a SALK T-DNA insertion located in the 5’ UTR of *PUB14*, which we named *pub14-1.* While a 5’ UTR insertion may have many different effects, we find that expression of the N-terminal portion of the *PUB14* gene is increased in the *pub14-1* mutant background (Figure 6 – Figure Supplement 1). We analyzed flowering time in the *pub14-1* mutant and compared it to the wild type. We observed that 1 cm bolting was delayed by 3.6 days in the *pub14-1* mutant and that it flowered with 4.5 more leaves on average (*p* < 0.0125; Figure 6), similar to the *PUB14* decoy population. Interestingly, we did not recapitulate the stem elongation defect observed in the *PUB14* decoy population, but do observe that anthesis is advanced by 1.6 days relative to wild type (*p* < 0.0125; Figure 6). The similarity in the phenotypes of the *PUB14* decoy population and the *pub14-1* mutant suggests that the *PUB14* is a *bona fide* regulator of flowering time, although additional experiments with a true knockout of *PUB14* would be beneficial for confirming its role in positive or negative regulation of flowering time.

**Figure 6.**
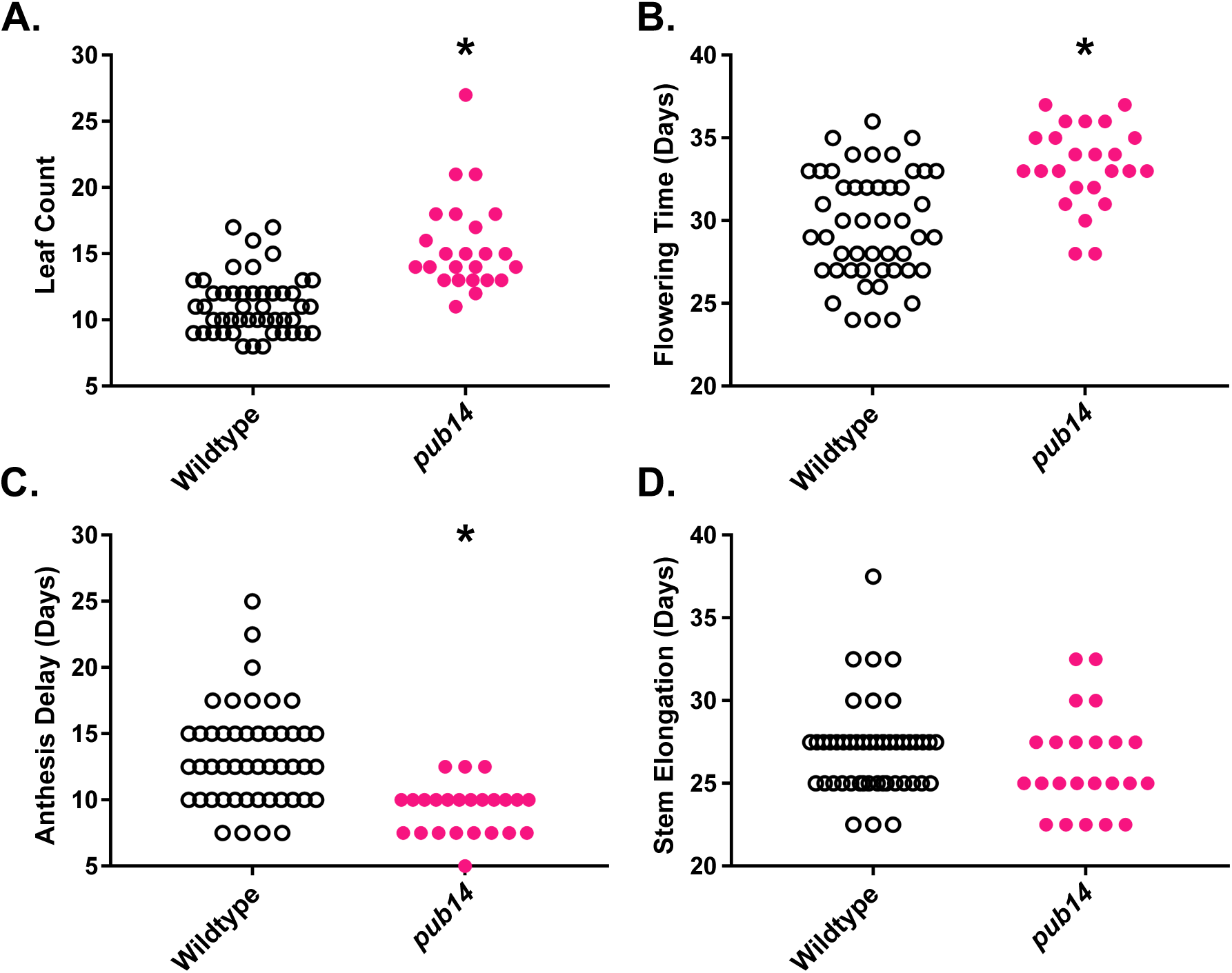
Flowering Time Analyses of *pub14-1* Mutants. A) Leaf number at 1 cm bolting. B) Age at 1 cm bolting. C) Anthesis delay. D) Elongation time. * represents a significant difference from wildtype with a Bonferroni-corrected *p* < 0.0125.

Reduction in *FT* expression levels is a hallmark of many late flowering mutants, although some mutants delay flowering independently of *FT* (Han et al., 2008; Leijten et al., 2018). In order to determine whether the *pub14-1* mutant delays flowering in an *FT-*dependent manner, we measured *FT* expression in wild type and *pub14-1* seedlings grown under long day conditions (Figure 7). In the wildtype plants, we observe patterns of *FT* expression corresponding to those observed previously (Song et al., 2012; Suárez-López et al., 2001; Wu et al., 2008). However, in the *pub14-1* mutant seedlings we observe a reduction of *FT* expression from ZT0 to ZT12. These results suggest that *PUB14* functions upstream of *FT* in flowering time regulation.

**Figure 7.**
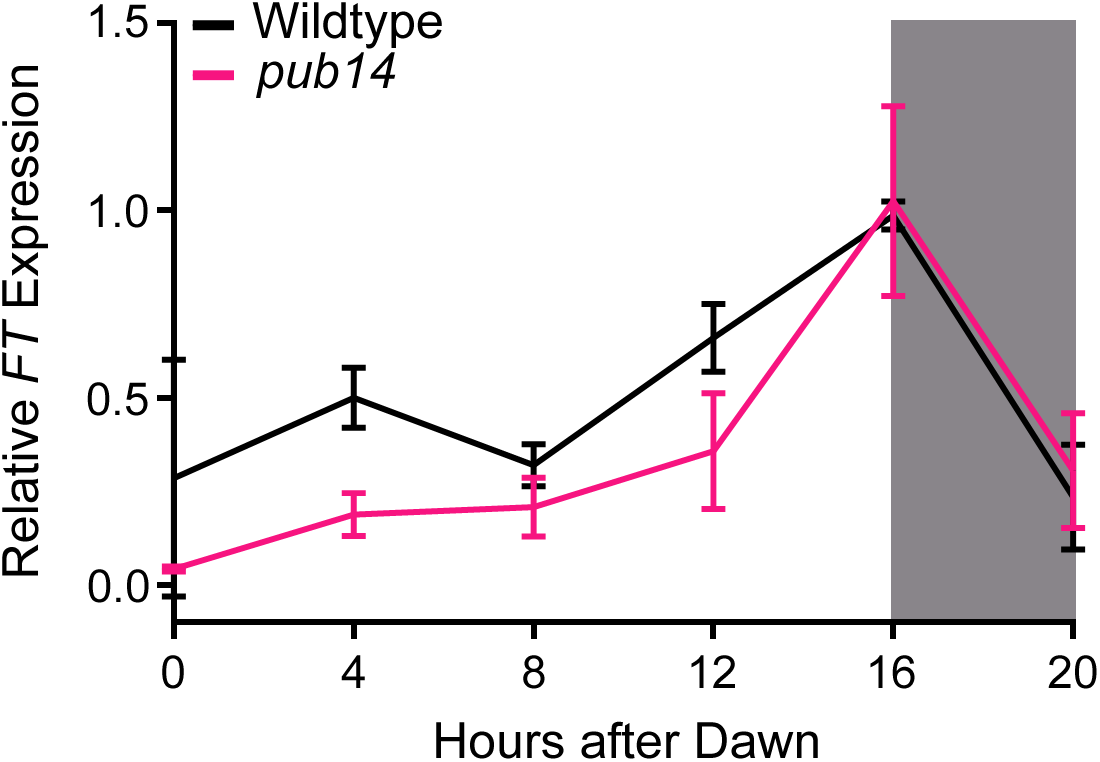
qRT-PCR of *FT* expression in *pub14-1* Mutants. *FT* expression was measured using quantitiative RT-PCR in wildtype or homozygous *pub14-1* mutants grown under long day (16 hours light/8 hours dark) conditions. Quantifications are the average of three biological replicates with error bars showing standard deviation.

### PUB14 Interacts with Flowering Time Regulators

Our data suggests that *PUB14* is a *bona fide* regulator of flowering time. However, it is unclear in which flowering time pathways *PUB14* functions. In order to better understand the biochemical function of PUB14, we performed immunoprecipitation followed by mass spectrometry on tissue expressing the 3XFLAG-6XHIS tagged PUB14 decoy and searched for flowering time regulators. We included tissue from plants expressing 3XFLAG-6XHIS tagged GFP as a control for proteins which bind to the tag, and wildtype parental plants as a control for proteins which bind to the beads. We identified four proteins with known function in flowering time regulation as potential interactors of PUB14 (Table 5.1). We identified peptides corresponding to SPLAYED (SYD), a SWI/SNF ATPase that represses flowering time under short day conditions, possibly by modulating activity of the floral activator LEAFY (LFY) (Wagner and Meyerowitz, 2002). We also identified peptides corresponding to SNW/SKI-Interacting Protein (SKIP), a component of the spliceosomal activating complex that represses flowering time through activating *FLC* (Cao et al., 2015; Cui et al., 2017). Similarly, we identify a potential interaction with ACTIN RELATED PROTEIN 6 (ARP6), a repressor of flowering that is required for *FLC* expression (Choi et al., 2005; Deal et al., 2005; Martin-Trillo et al., 2006). Finally, we identify *TOPLESS* (*TPL*), a protein that acts as a weak repressor of flowering time, potentially by forming a complex with CO to repress *FT* activation (Causier et al., 2012; Graeff et al., 2016). While additional work is required to verify these interactions and test whether they are ubiquitylation targets of PUB14, the identification of flowering time repressors as putative interacting partners of PUB14 may explain the late flowering phenotype we observe in the *pub14-1* mutant and *PUB14* decoy-expressing plants.

We have previously observed that E3 ligases interact with close homologs (Feke et al., 2019; Lee and Feke et al., 2018). Thus, we searched our IP-MS data for other U-box genes that interact with PUB14. We did not identify peptides corresponding to PUB13, the closest homolog of PUB14. However, we did identify peptides corresponding to two other close homologs of PUB14, PUB12 (E-value 3×10^-163^) and PUB10 (E-value 8×10^-142^). Interestingly, we do not identify peptides corresponding to the other members of this small subfamily, PUB15 (E–value 6×10^-134^) and PUB11 (E–value 4×10^-142^). While the importance of these interactions has not been verified, this data suggests that interaction between homologs is a common feature of E3 ligase complexes, and that PUB10 and PUB12 may also be involved in flowering time regulation.

### *MAC3A* and *MAC3B* Regulate Flowering Time in a Partially Redundant Manner

Expression of the *MAC3A* decoy leads to the greatest magnitude change that we observed in our screen (2.9 more leaves than wild type, Figure 1). *MAC3A* and *MAC3B* can act as fully or partially redundant regulators of processes controlled by the plant spliceosomal activating complex (Feke et al., 2019; Jia et al., 2017; Li et al., 2018; Monaghan et al., 2009). It was previously noted that *MAC3A* and *MAC3B* could regulate flowering time (Monaghan et al., 2009). We have established genetic tools to further investigate the genetic interaction of *MAC3A* and *MAC3B* in flowering time. We grew the single and double mutants in an inductive photoperiod and measured our four flowering time parameters (Figure 8). We observe a statistically significant difference in leaf number at 1 cm bolting between all three mutant backgrounds and the wildtype, but observe no difference between the mutant backgrounds (5.3, 4.0, and 5.1 more leaves than wild type in the *mac3a*, *mac3b*, and *mac3a/mac3b* mutants, respectively; Figure 8a). Genetically this indicates that these two genes are in series or function together for this aspect of flowering time control. For the number of days to 1 cm bolting (flowering time; Figure 8b), we observe a statistical difference from wildtype in all three backgrounds, with the double mutant being the most delayed (10.7 days), the *mac3b* mutant being the least delayed (5.4 days) and the *mac3a* mutant having an intermediate delay in flowering (7.1 days). The increase in severity of the double mutant indicates that *MAC3A* and *MAC3B* can act redundantly for 1 cm bolting. The anthesis delay is shorter in the singe mutants than in the wild type (1.5 and 1.3 days for the *mac3a* and *mac3b* mutants, respectively; Figure 8c), but is indistinguishable from wildtype in the *mac3a/mac3b* double mutant. The stem elongation time is also shorter in the *mac3a* single mutant when compared to the wild type (0.8 days; Figure 8d), but longer in the *mac3a/mac3b* double mutant (0.8 days). We observe no statistical difference in stem elongation between the *mac3b* single mutant and the wild type. What is clear from this data is that *MAC3A* and *MAC3B* are necessary for the plant to properly time developmental transitions. This experiment also confirms what has previously been seen that *MAC3A* and *MAC3B* can act partially redundantly and possibly together to control important biological processes.

**Figure 8.**
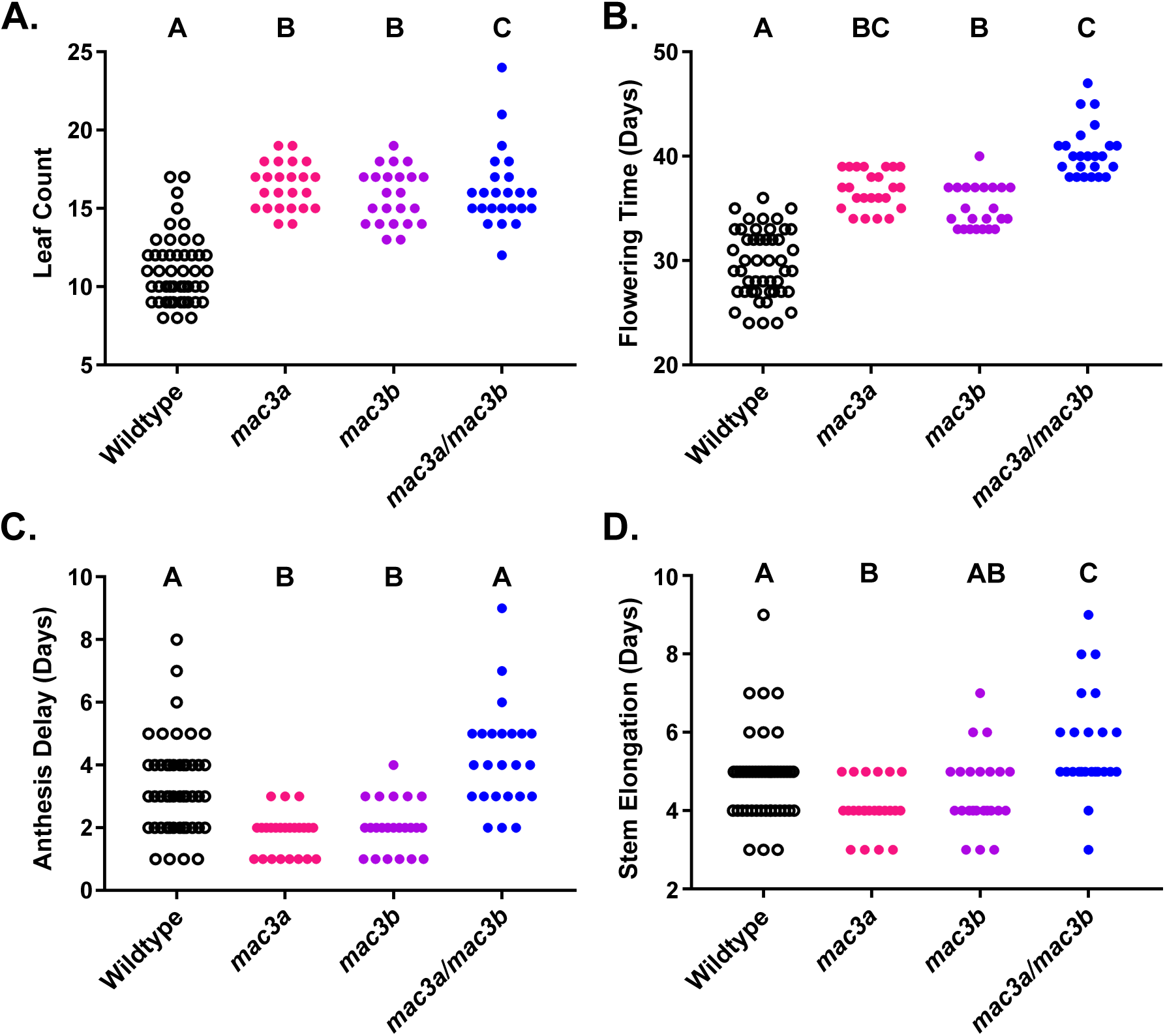
Flowering Time Analyses of *mac3A*, *mac3B*, and *mac3A/mac3B* Mutants. A) Leaf number at 1 cm bolting. B) Age at 1 cm bolting. C) Anthesis delay. D) Elongation time. Letters represent statistical groups as defined by a Kruskal-Wallis test with a post-hoc Dunn’s multiple comparisons test, with statistical differnece defined as *p* < 0.05.

In order to determine whether the flowering time delays we observe in the *mac3a*, *mac3b*, and *mac3a/mac3b* mutants are due to an *FT-*dependent or –independent process, we measured *FT* expression in these plants using qRT-PCR. We observe a decrease in *FT* expression in all three mutant backgrounds. The decrease in *FT* expression is maximal from ZT0-ZT8 in the single *mac3a* and *mac3b* mutants, although *mac3b* also has decreased *FT* expression at the ZT16 peak. In contrast, the maximal decrease in *FT* levels in the *mac3a/mac3b* double mutant occurs at ZT16. The *mac3a/mac3b* double mutant shows an even greater decrease in *FT* expression at the peak, which may explain the greater delay in flowering time in this background. These results strongly indicate that *MAC3A* and *MAC3B* are functioning upstream of *FT* to control flowering.

### Ubiquitylation Dependency of MAC3B on Flowering Time Control

We have shown that *MAC3A* and *MAC3B* act as partially redundant regulators of flowering time, and we have shown that *MAC3A* and *MAC3B* can form a heterodimer complex in plants in the absence of the U-box domain (Feke et al., 2019). In order to test the role of MAC3A/MAC3B dimerization in flowering time regulation, we created a *MAC3B* decoy construct which consists of only the annotated WD40 repeats and is missing the canonical coiled-coil domain required for oligomerization (Grote et al., 2010; Ohi et al., 2005). We were unable to generate similar constructs for *MAC3A* due to unknown technical constraints, as described previously (Feke et al., 2019). We conducted the flowering time assays with plants expressing the *MAC3B* WD construct, and included *MAC3B* decoy plants as control. The transgenic plants expressing the *MAC3B* WD delays 1 cm bolting by 4.5 days (*p* = 3.8×10^-12^), compared to the decoy expressing plants which delayed flowering by 1.45 days (*p* = 0.016) (Figure 10). Interestingly, this was not the case in our other flowering time metrics, as leaf number, anthesis delay, and stem elongation period was identical to the wildtype in the *MAC3B* WD population. The *MAC3B* decoy had similar trends in anthesis delay and elongation period defects that we observed in our initial screen, delaying flowering by 0.95 (*p* = 3×10^-3^) and 0.88 days (*p* = 0.039) respectively, but did not have any effects on leaf number. Taken together, this suggests that the anthesis delay and stem elongation defects that we observe in the *MAC3B* decoy is dependent on its ability to form protein dimers, while the delayed 1 cm bolting is a dominant negative effect.

**Figure 9.**
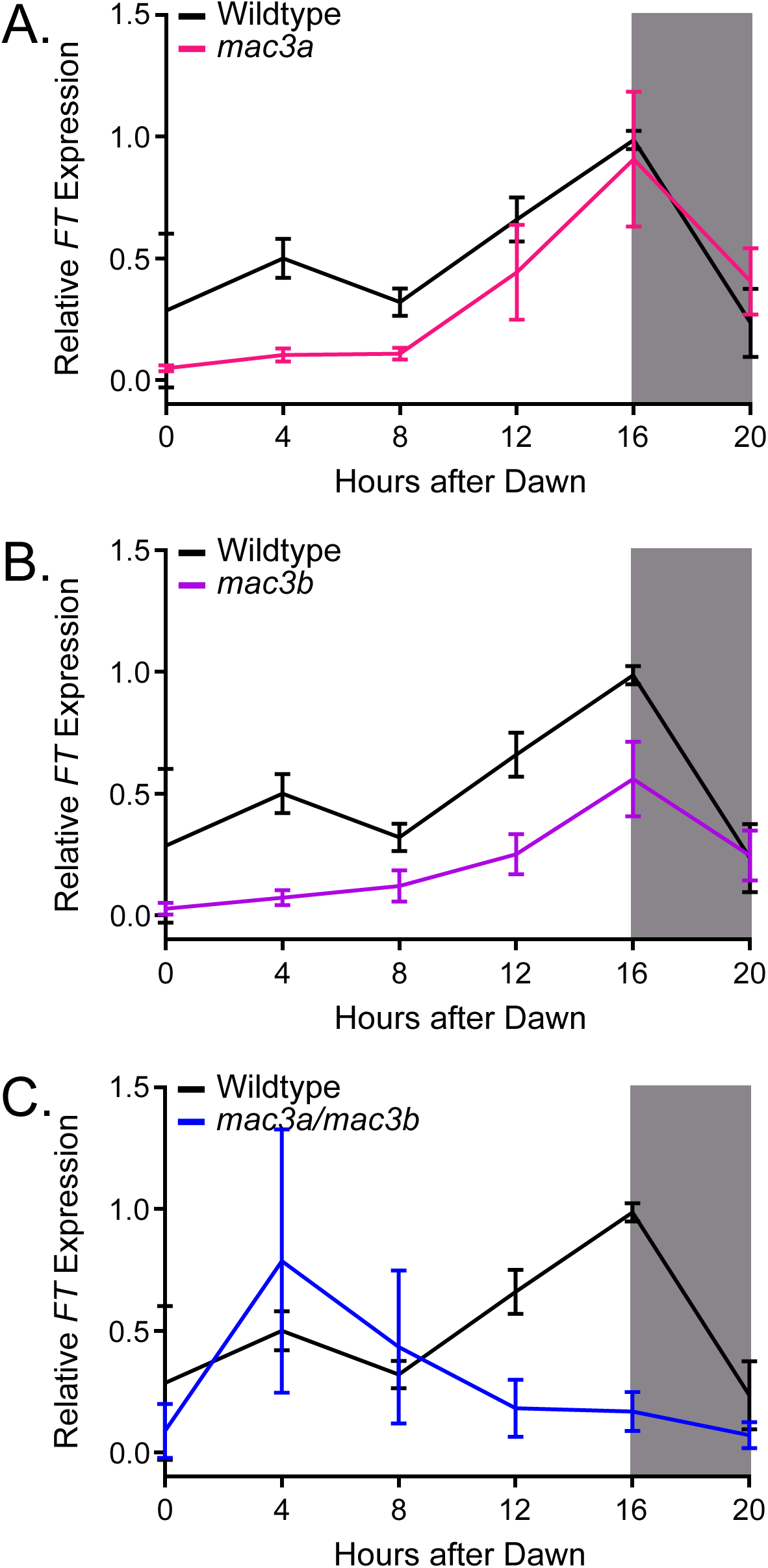
qRT-PCR of *FT* expression in *mac3A, mac3B, and mac3A/mac3B* Mutants. *FT* expression was measured using quantitiative RT-PCR in wildtype or homozygous A) *mac3A* B) *mac3B* and C) *mac3a/mac3B* mutants grown under long day (16 hours light/8 hours dark) conditions. Quantifications are the average of three biological replicates with error bars showing standard deviation.

**Figure 10.**
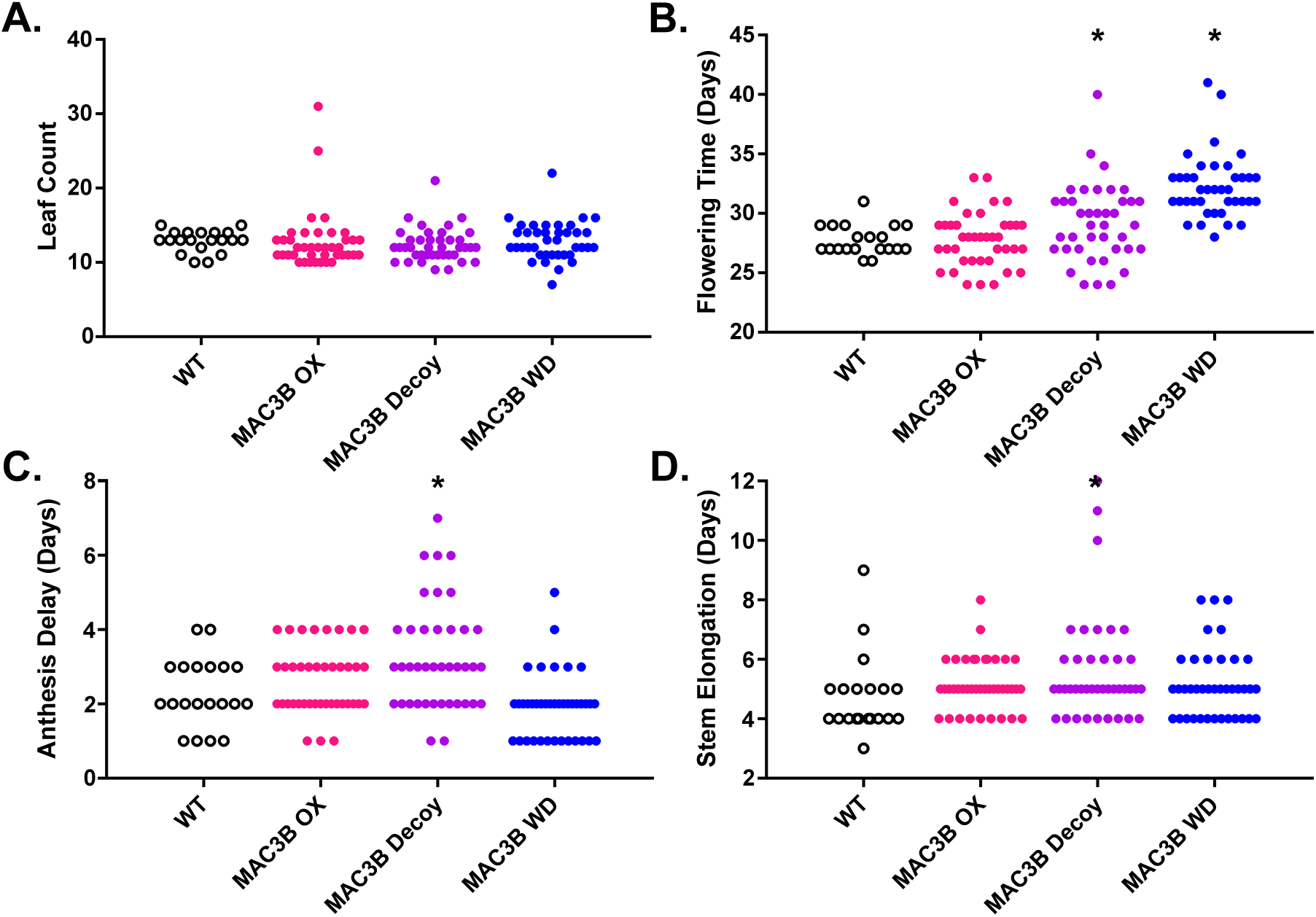
Flowering Time Analyses of *MAC3B* Overexpression Constructs. Flowering time was measured in T1 *MAC3B* full length (OX), *MAC3B* decoy, and *MAC3B* WD insertion plants. A) Age at 1 cm inflorescence. B) Leaf number at 1 cm inflorescence. C) Anthesis delay. D) Stem elongation time. Brackets define individual groups used for statistical testing against the wildtype control. * represents a significant difference from wildtype with a Bonferroni-corrected *p* < 0.017.

We have previously reported that precise regulation of *MAC3B* expression is essential for maintaining periodicity in the circadian clock (Feke et al., 2019). However, it is unclear if flowering time is also sensitive to *MAC3B* expression levels. Thus, we overexpressed the full-length *MAC3B* and assayed its effects on flowering time. Interestingly, we do not observe any alteration in flowering time in the full-length *MAC3B* overexpression plants (Figure 10). This suggests that the role of *MAC3B* in the regulation of flowering time relies on its ability to ubiquitylate substrates and not on precise regulation of *MAC3B* expression levels.

## DISCUSSION

We have previously demonstrated the utility of the decoy technique to overcome redundancy and identify E3 ligases which regulate the circadian clock (Feke et al., 2019; Lee et al., 2018). Here, we demonstrate that the decoy technique is capable of identifying E3 ligases involved in developmental processes, specifically flowering time regulation. We were able to identify five major candidates and ten minor candidates for flowering time regulators, and performed follow-up experiments to validate two of the major candidates. While one of these candidates, *MAC3A,* and its homolog, *MAC3B*, have been suggested to control flowering time previously (Monaghan et al., 2009), we directly demonstrate flowering defects in single and double mutants and reveal the complicated partial redundancy between the two genes. This study and our previous study on the role of *MAC3A* and *MAC3B* in clock function, will likely help to clarify the genetic roles of *MAC3A* and *MAC3B* in the myriad biological processes that they control (Feke et al., 2019; Jia et al., 2017; Li et al., 2018; Monaghan et al., 2009). Furthermore, a second candidate revealed by our screen, *PUB14*, had not been identified for its role in any biological process previously, but has sequence similarity to the characterized flowering time regulator *PUB13* (Li et al., 2012b, 2012a; Zhou et al., 2015).

### *PUB14* Is a Novel Flowering Time Regulator

*PUB14* was the only candidate flowering time regulator found in our screen that affected both leaf number and 1 cm bolting age, two hallmarks of flowering time (Koornneef et al., 1991). Although the biochemical structure of the PUB14 protein has been studied (Andersen et al., 2004), to our knowledge no phenotypes have previously been associated with mutations in this gene. Here, we validate *PUB14* as a regulator of flowering time. Both *PUB14* decoys and the *pub14-1* T-DNA insertion mutant have delayed 1 cm bolting and increased leaf number, Furthermore, *FT* expression is reduced in the *pub14-1* mutant. As we find that the *pub14-1* mutant has increased expression of a portion of the *PUB14,* the similarity in phenotype between the *PUB14* decoy and the *pub14-1* mutant suggests that the decoy could be acting in a dominant positive manner in this case, rather than a dominant negative. We have previously observed dominant positive effects with the other decoys in relation to the regulation of the circadian clock and flowering time (Feke et al., 2019; Lee and Feke et al., 2018).

The closest homolog of *PUB14* is *PUB13*, which has previously been implicated in stress responses and the control of flowering time (Li et al., 2012b, 2012a; Zhou et al., 2015). Mutants of *PUB13* have accelerated flowering time under long day growth conditions, suggesting that *PUB13* acts as a repressor of photoperiodic flowering (Li et al., 2012a, 2012b; Zhou et al., 2015). *pub13* mutants also have elevated levels of the defense hormone salicylic acid (SA), suggesting that *PUB13* is a negative regulator of immunity (Li et al., 2012a). SA activates flowering (Li et al., 2012a; Martínez et al., 2004), indicating PUB13 is a repressor of flowering time and immunity through negatively regulating SA levels. Correspondingly, the advanced flowering time in the *pub13* mutant is dependent on SA (Li et al., 2012a; Zhou et al., 2015). We were unable to recapitulate the *pub13* mutant flowering phenotype with the *PUB13* decoy, although we do see a trend towards advanced flowering that does not reach our statistical cutoff (Figure 2). However, the high protein sequence similarity in the substrate recognition domains of PUB14 and PUB13 (83% similar) and the similarity of their functions suggests that these two homologous genes may share the same targets. In addition to its potential role in stress-regulated flowering, we also identify a suite of potential flowering time regulators that interact with *PUB14* in our IP-MS experiments. Direct interaction studies such as yeast-two-hybrid or co-immunoprecipitation are required to verify that these putative substrates interact with PUB14. Further genetic and molecular studies can determine whether they are targets or regulatory partners of PUB14 (Lee et al., 2018, 2019).

### *MAC3A* Regulates Flowering Time

*MAC3A*, (also known as *MOS4-ASSOCIATED COMPLEX 3A* (*MAC3A*) or *PRE-mRNA PROCESSING FACTOR 19 A* (*PRP19A*)) and its close homolog *MAC3B* (*MAC3B/PRP19B*) are the core component of a large, multi-functional protein complex known as the Nineteen Complex (NTC) (Monaghan et al., 2009). In plants, the NTC, and *MAC3A* and *MAC3B* in particular, has been implicated in splicing, miRNA biogenesis, immunity, and the circadian clock (Feke et al., 2019; Jia et al., 2017; Li et al., 2018; Monaghan et al., 2009). For this reason, we don’t believe that MAC3A and MAC3B would be interacting with and ubiquitylating proteins that regulate flowering, but would rather alter processes through splicing or other NTC processes. Interestingly, another component of the NTC, SKIP, has been implicated in flowering time regulation previously (Cao et al., 2015; Cui et al., 2017). Here, we identified *MAC3A* as the U-box decoy with the greatest magnitude effect on flowering time, and validated that *MAC3A* and *MAC3B* are *bona fide* regulators of flowering time. We do observe different phenotypes between the *MAC3A* decoy, *MAC3B* decoy, and *mac3a/mac3b* mutants, which suggests a complex relationship with flowering time. However, it is clear that both genes are essential for proper flowering time control.

The precise methods through which *MAC3A* and *MAC3B* alter flowering time are not yet understood and likely multi-factorial. *MAC3A* and *MAC3B* are involved in regulation of splicing, miRNA biogenesis, immunity, and the circadian clock (Feke et al., 2019; Jia et al., 2017; Li et al., 2018; Monaghan et al., 2009). Interestingly, all of these are involved in the regulation of flowering time (Chen, 2004; Cui et al., 2017; Gil et al., 2017; Imaizumi et al., 2003; Lyons et al., 2015; Wu et al., 2009; Wu and Poethig, 2006; Yamaguchi et al., 2009; Yanovsky and Kay, 2002; Yant et al., 2010). Alterations in the circadian clock lead to defects in photoperiodic flowering time, similar to what we observe in the *mac3a/mac3b* double mutant (Nakamichi et al., 2007). Likewise, increased resistance to pathogens, like what is observed in the *mac3a/mac3b* double mutant, is positively correlated with a delay in flowering time (Korves and Bergelson, 2003; Lyons et al., 2015; Monaghan et al., 2009). miRNAs play an essential role in the regulation of flowering time through the aging pathway, with miRNAs having both activating and repressive activity within this pathway (Chen, 2004; Wu et al., 2009; Wu and Poethig, 2006; Yamaguchi et al., 2009; Yant et al., 2010). However, interpretation of the relationship between *MAC3A* and *MAC3B* and this pathway is complicated by the fact that both the repressive and activating miRNAs are likely affected by these genes (Jia et al., 2017; Li et al., 2018). Finally, splicing also plays a role in the regulation of flowering, as both the photoperiodic floral activator *CO* and ambient temperature floral repressor *FLM* are alternatively spliced (Gil et al., 2017; Lee et al., 2013; Posé et al., 2013). In addition, our results suggest that the ubiquitylation activity of MAC3B is essential for its ability to regulate flowering time. In our truncation studies, we observed an anti-correlation between the presence of the U-box domain and proper regulation of flowering time, with no effect on flowering time observed in plants overexpressing full-length MAC3B and the largest impact on flowering time in plants expressing the putative substrate interaction domain alone. Future investigation into the relationships between the diverse functions of *MAC3A* and *MAC3B* and flowering time will likely prove fruitful.

### Additional Flowering Time Candidates Connect Stress to Flowering Time

Stress is a well-known regulator of flowering time in Arabidopsis (Takeno, 2016). Correspondingly, all of our remaining high-priority candidate floral regulators have established roles in stress responses. *PUB26* is a negative regulator of immunity, and *pub26* mutants exhibit elevated levels of immunity (Wang et al., 2018). As resistance to pathogens and delayed flowering are positively correlated (Lyons et al., 2015), we would expect that that flowering time would be delayed in these mutants, in concordance with our observations of flowering time in the *PUB26* decoy population. Like biotic stresses, abiotic stresses such as salt stress can delay flowering time (Kim et al., 2007). In accordance with this, we observe delayed flowering with the *PUB31* decoy, which leads to mild sensitivity to salt stress when mutated (Zhang et al., 2017). In contrast, the correlation between the known stress phenotypes of *PUB61*, also known as *CARBOXYL TERMINUS OF HSC70-INTERACTING PROTEIN* (*CHIP*), is less easily interpretable. We observe early flowering with the *CHIP* decoy. CHIP was previously identified to alter sensitivity to heat, cold, salt and ABA, but the system is complicated because mutants and overexpression lines are both sensitive to these stresses (Luo et al., 2006; Wei et al., 2015; Zhou et al., 2014). In this case, use of the decoy may help to untangle the gmcomplex relationships between *CHIP*, stress, and flowering time.

### Flowering Time Metrics are Not Equivalent

In our study, we investigated four different metrics of flowering time: the leaf number at 1 cm bolting, the age at 1 cm bolting, the anthesis delay, and the amount of time that it takes for a stem to elongate from 1 cm to 10 cm, a proxy for the stem elongation rate. By investigating these metrics, we were able to get a more comprehensive picture of floral development for all of our decoy and mutant populations. By analyzing this data, it is clear that these metrics are not interchangeable with one another. The majority of the decoy populations screened in this study which had a defect in flowering time only affected one of the metrics. Furthermore, as exhibited by the complex genetic interactions we observe in the *mac3a*, *mac3b*, and *mac3a/mac3b* mutants, genes that similarly affect one flowering time metric may affect other flowering time metrics differently. Flowering is a complex process that includes many steps from the initiation of the floral meristem to finally anthesis. Our study demonstrates that there can be different genetic systems involved in the transitions between each one of these smaller steps in what we know as flowering. Further work will be required to untangle the complex relationships between these various aspects of floral development timing.

### Conclusions

A multitude of factors, ranging from light conditions and temperature to the effects of stress, contribute to the regulation of flowering time. We only selected one condition, the floral inductive long day condition, to perform our screen, and due to the labor intensiveness of this screen, we chose to only investigate the U-box library. Despite using these limited conditions, we were able to identify five novel regulators of flowering time, and validated two by mutant analysis. This demonstrates the likely magnitude of undiscovered flowering time regulators within the E3 ligases as a whole, and demonstrates the necessity of targeted, dominant-negative screens to characterize members of these complex gene classes. Our experimental procedures and results provide a model for future studies of the roles of E3 ligases in flowering time and other developmental processes, and solidify the usefulness of the decoy technique as a screening platform for identifying plant E3 ligase function.

## MATERIALS AND METHODS

### Phenotypic Screening

The construction of the decoy library, the *MAC3B-OX,* and the *MAC3B-WD* was described previously (Feke et al., 2019). Control *pCCA1∷Luciferase* and decoy seeds were surface sterilized in 70% ethanol and 0.01% Triton X-100 for 20 minutes prior to being sown on ½ MS plates (2.15 g/L Murashige and Skoog medium, pH 5.7, Cassion Laboratories, cat#MSP01 and 0.8% bacteriological agar, AmericanBio cat# AB01185) with or without appropriate antibiotics (15 μg/mL ammonium glufosinate (Santa Cruz Biotechnology, cat# 77182-82-2) for vectors pB7-HFN and pB7-HFC, or 50 μg/mL kanamycin sulfate (AmericanBio) for pK7-HFN). Seeds were stratified for two days at 4 °C, transferred to 12 hours light/12 hours dark conditions for seven days, then to constant light conditions for 7 days in order to do screening for circadian clock studies shown in Feke *et al.* 2019. Seedlings were then transferred to soil (Fafard II) and grown at 22 °C in inductive 16 hours light/8 hours dark conditions with a light fluence rate of 135 μmol m-2. Plants were monitored daily for flowering status, recording the dates upon which each individual reached 1 cm inflorescence height, 10 cm inflorescence height, and the first occurrence of anthesis. Additionally, leaf number at 1 cm inflorescence height was recorded.

Homozygous *pub14-1*, *mac3a*, *mac3b*, and *mac3a/mac3b* mutant seeds were surface sterilized, sown on ½ MS plates without antibiotics as described above. Seeds were stratified for 3 days at 4°C, then transferred to 12 hours light/ 12 hours dark conditions for two weeks prior to transfer to soil and growth under inductive conditions as described above. Plants were monitored daily for flowering status as described above.

### Data Normalization and Statistical Analysis

As the age at anthesis depends on the initiation of flowering, we used anthesis delay as a measurement of anthesis. Anthesis delay was calculated by taking the age at anthesis and subtracting the age at 1 cm inflorescence height. Similarly, the age at 10 cm inflorescence height depends on the initiation of flowering. Thus we calculated the stem elongation period by subtracting the age at 1 cm inflorescence height from the age at 10 cm inflorescence height. These modified metrics were used for all analyses.

To allow for comparison across independent experiments, data was normalized to the individual wildtype control performed concurrently. The average value of the wildtype control plants was calculated for every experiment, then this average was subtracted from the value of each individual T1 insertion or control wildtype plant done concurrently. This normalized value was used for statistical analyses.

Welch’s t-test was used to compare each normalized T1 insertion plant population or subpopulation to the population of all normalized control plants. In order to decrease the number of false positives caused by multiple testing, we utilized a Bonferroni corrected α as the p-value threshold. The α applied differs between experiments, and is noted throughout.

### Measurement of Gene Expression in U-box mutants

Homozygous *mac3a/mac3b* mutant plants in the Col-0 background were generated previously (Monaghan et al., 2009). Col-0, *pub14-1*, *mac3a*, *mac3b*, and *mac3a/mac3b* seeds were stratified on ½ MS plates at 4 °C for two days prior to growth in 16 hr light/8 hr dark conditions at a fluence rate of 130 μmol m^−2^ s^−1^ at 22 °C. 10-day old seedlings were collected in triplicate every four hours for one day starting at ZT0 and snap-frozen using liquid nitrogen, then ground using the Mixer Mill MM400 system (Retsch). Total RNA was extracted from ground seedlings using the RNeasy Plant Mini Kit and treated with RNase-Free DNase (Qiagen, cat#74904 and 79254) following the manufacturer’s protocols. cDNA was prepared from 1 μg total RNA using iScript^TM^ Reverse Transcription Supermix (Bio-Rad, cat#1708841), then diluted 10-fold and used directly as the template for quantitative real-time RT-PCR (qRT-PCR). The qRT-PCR was performed using 3.5 μl of diluted cDNA and 5.5 μM primers listed in Table 2 (C.-M. Lee and Thomashow, 2012; Wu et al., 2008)using iTaq^TM^ Universal SYBR^®^ Green Supermix (Bio-Rad, cat# 1725121) with the CFX 384 Touch^TM^ Real-Time PCR Detection System (Bio-RAD). The qRT-PCR began with a denaturation step of 95°C for 3 min, followed by 45 cycles of denaturation at 95°C for 15 sec, and primer annealing at 53°C for 15s. Relative expression was determined by the comparative C_T_ method using *IPP2* (*AT3G02780*) as an internal control. The relative expression levels represent the mean values of 2^-ΔΔCT^ from three biological replicates, where ΔCT = C_T_ of *FT* – C_T_ of *IPP2* and the reference is Col-0 replicate #1. When measuring *FT* expression, the time point of peak expression (ZT16) was used as the reference point.

**Table 1.**
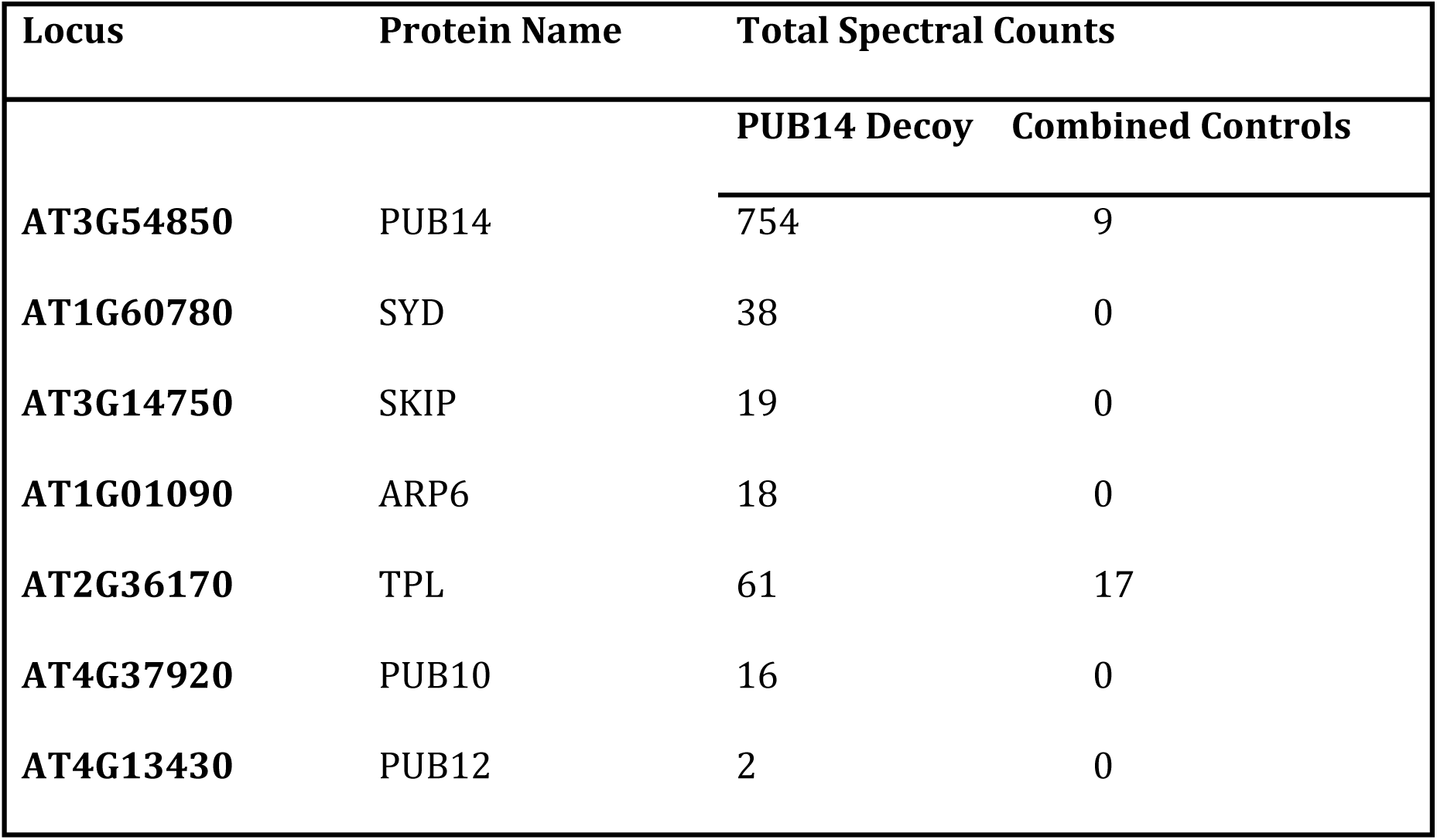
Selected IP-MS Results from the PUB14 Decoy. PUB14 decoy peptide hits are from one IP-MS experiment using the PUB14 decoy as the bait. Combined control peptide hits are summed from the independent control experiments of wildtype Col-0 and *35S::His-FLAG-GFP* expressing plants.

| Locus | Protein Name | Total Spectral Counts |  |
| --- | --- | --- | --- |
|  |  | PUB14 Decoy | Combined Controls |
| AT3G54850 | PUB14 | 754 | 9 |
| AT1G60780 | SYD | 38 | 0 |
| AT3G14750 | SKIP | 19 | 0 |
| AT1G01090 | ARP6 | 18 | 0 |
| AT2G36170 | TPL | 61 | 17 |
| AT4G37920 | PUB10 | 16 | 0 |
| AT4G13430 | PUB12 | 2 | 0 |

**Table 2.**
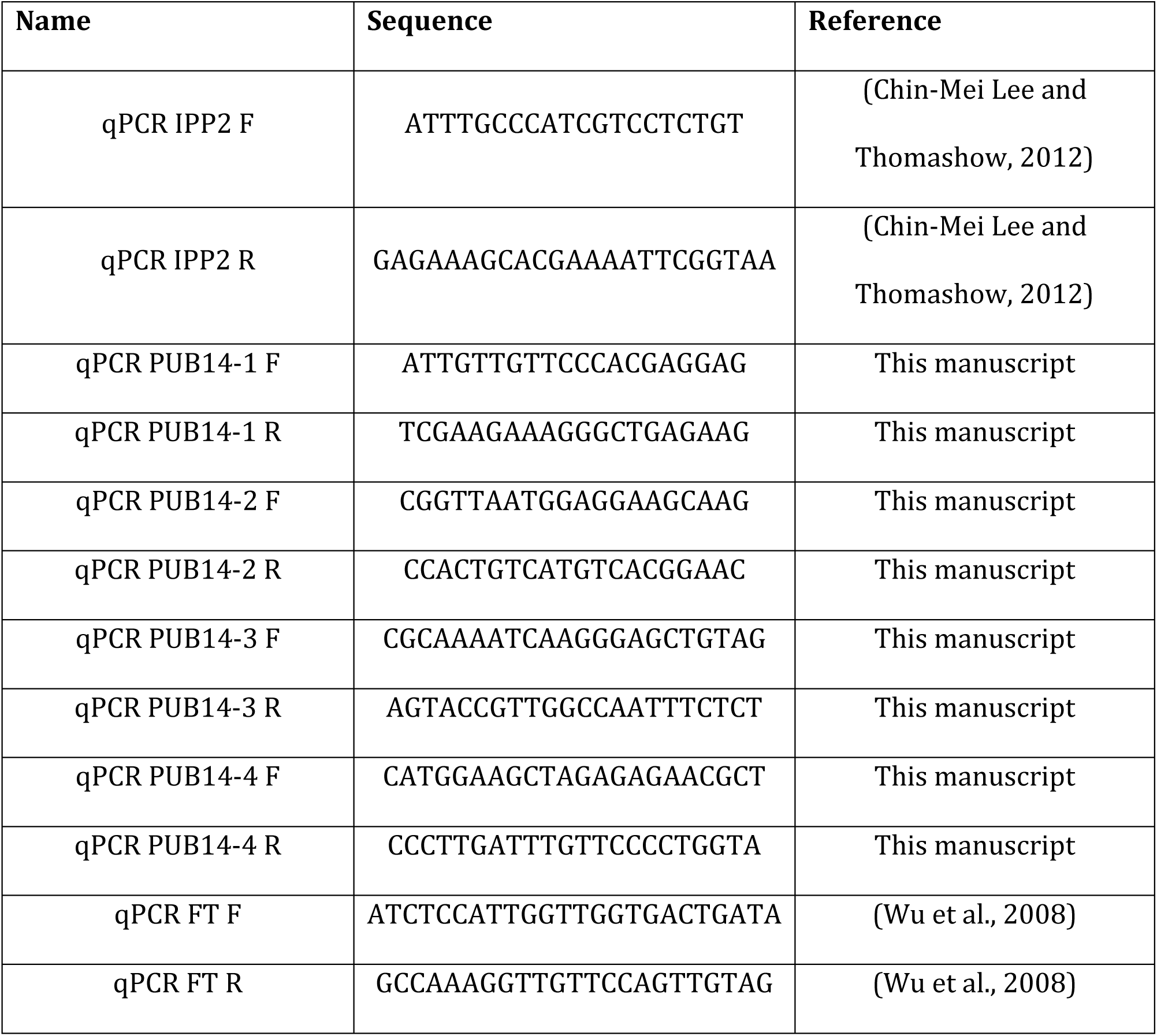
Primers used in this study.

### Immunoprecipitation and Mass Spectrometry of PUB14 Decoy plants

Individual T1 *pB7-HFN-PUB14* transgenic plants in a Col-0 background and control Col-0 and *pB7-HFC-GFP* were grown as described for phenotype analysis. Seven-day old seedlings were transferred to soil and grown under 16 hours light/8 hours dark at 22 °C for 2-3 weeks. Prior to harvest, plants were entrained to 12 hours light/12 hours dark at 22 °C for 1 week. Approximately 40 mature leaves from each background was collected and flash frozen in liquid nitrogen, such that each sample was a mixture of leaves from multiple individuals to reduce the effects of expression level fluctuations. Tissue samples were ground in liquid nitrogen using the Mixer Mill MM400 system (Retsch). Immunoprecipitation was performed as described previously (Huang et al., 2016a, 2016b; Lu et al., 2010). Briefly, protein from 2 mL tissue powder was extracted in SII buffer (100 mM sodium phosphate pH 8.0, 150 mM NaCl, 5 mM EDTA, 0.1% Triton X-100) with cOmplete™ EDTA-free Protease Inhibitor Cocktail (Roche, cat# 11873580001), 1 mM phenylmethylsµlfonyl fluoride (PMSF), and PhosSTOP tablet (Roche, cat# 04906845001) by sonification. Anti-FLAG antibodies were cross-linked to Dynabeads^®^ M-270 Epoxy (Thermo Fisher Scientific, cat# 14311D) for immunoprecipitation. Immunoprecipitation was performed by incubation of protein extracts with beads for 1 hour at 4 °C on a rocker. Beads were washed with SII buffer three times, then twice in F2H buffer (100 mM sodium phosphate pH 8.0, 150 mM NaCl, 0.1% Triton X-100). Beads were eluted twice at 4 °C and twice at 30 °C in F2H buffer with 100 μg/mL FLAG peptide, then incubated with TALON magnetic beads (Clontech, cat# 35636) for 20 min at 4 °C, then washed twice in F2H buffer and three times in 25 mM Ammonium Bicarbonate. Samples were subjected to trypsin digestion (0.5 µg, Promega, cat# V5113) at 37 °C overnight, then vacuum dried using a SpeedVac before being dissolved in 5% formic acid/0.1% trifluoroacetic acid (TFA). Protein concentration was determined by nanodrop measurement (A260/A280)(Thermo Scientific Nanodrop 2000 UV-Vis Spectrophotometer). An aliquot of each sample was further diluted with 0.1% TFA to 0.1µg/µl and 0.5µg was injected for LC-MS/MS analysis at the Keck MS & Proteomics Resource Laboratory at Yale University.

LC-MS/MS analysis was performed on a Thermo Scientific Orbitrap Elite mass spectrometer equipped with a Waters nanoACQUITY UPLC system utilizing a binary solvent system (Buffer A: 0.1% formic acid; Buffer B: 0.1% formic acid in acetonitrile). Trapping was performed at 5µl/min, 97% Buffer A for 3 min using a Waters Symmetry® C18 180µm x 20mm trap column. Peptides were separated using an ACQUITY UPLC PST (BEH) C18 nanoACQUITY Column 1.7 µm, 75 µm x 250 mm (37°C) and eluted at 300 nL/min with the following gradient: 3% buffer B at initial conditions; 5% B at 3 minutes; 35% B at 140 minutes; 50% B at 155 minutes; 85% B at 160-165 min; then returned to initial conditions at 166 minutes. MS were acquired in the Orbitrap in profile mode over the 300-1,700 m/z range using 1 microscan, 30,000 resolution, AGC target of 1E6, and a full max ion time of 50 ms. Up to 15 MS/MS were collected per MS scan using collision induced dissociation (CID) on species with an intensity threshold of 5,000 and charge states 2 and above. Data dependent MS/MS were acquired in centroid mode in the ion trap using 1 microscan, AGC target of 2E4, full max IT of 100 ms, 2.0 m/z isolation window, and normalized collision energy of 35. Dynamic exclusion was enabled with a repeat count of 1, repeat duration of 30s, exclusion list size of 500, and exclusion duration of 60s.

The MS/MS spectra were searched by the Keck MS & Proteomics Resource Laboratory at Yale University using MASCOT (Perkins et al., 1999). Data was searched against the SwissProt_2015_11.fasta *Arabidopsis thaliana* database with oxidation set as a variable modification. The peptide mass tolerance was set to 10 ppm, the fragment mass tolerance to 0.5 Da, and the maximum number of allowable missed cleavages was set to 2.

## ACKNOWLEDGEMENTS

We would like to thank Christopher Adamchek, Cathy Chamberlin, Suyuna Eng Ren, Brandon Williams, Milan Sandhu, Annie Jin, and Skylar McDermott for their technical support. We would also like to thank Adam Saffer for his critical reading of the manuscript. Additionally, we would like to thank Chris Bolick, Eileen Williams, and the staff at Marsh Botanical Gardens for their support in maintaining plant growth spaces. We would also like to thank Sandra Pariseau and Denise George for their administrative support. Finally, we would also like to thank the Keck Proteomics Facility at Yale for their assistance with proteomics, and Dr. Xin Li for providing the *mac3a* and *mac3a/mac3b* double mutants.

A.F. and J.M.G. designed the experiments. A.F. performed the experiments and experimental analyses. A.F., J.H, and W.L. were involved in the generation of the U-box decoy library. A.F. and J.M.G. wrote the manuscript.

This work was supported by the National Science Foundation (EAGER #1548538) and the National Institutes of Health (R35 GM128670) to J.M.G; by the Forest B.H. and Elizabeth D.W. Brown Fund Fellowship to W.L.; and by the National Institutes of Health (T32 GM007499), the Gruber Foundation, and the National Science Foundation (GRFP DGE-1122492) to A.F.

## COMPETING INTERESTS

The authors declare no competing interests.

## FIGURE LEGENDS

**Figure 1 – Figure Supplement 1.**
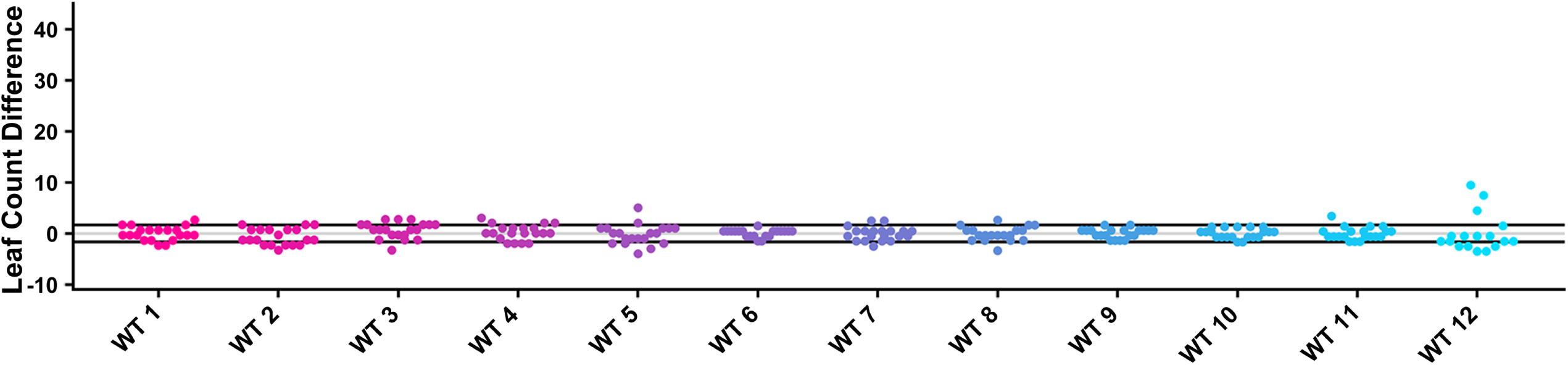
Leaf Count Distributions of Control Plants. Values presented are the difference between the leaf count at 1 cm inflorescence of the individual control plant and the average leaf count of the control in the accompanying experiment. The grey line is at the average control value and the black lines are at +/- the standard deviation of the control plants.

**Figure 2 – Figure Supplement 1.**
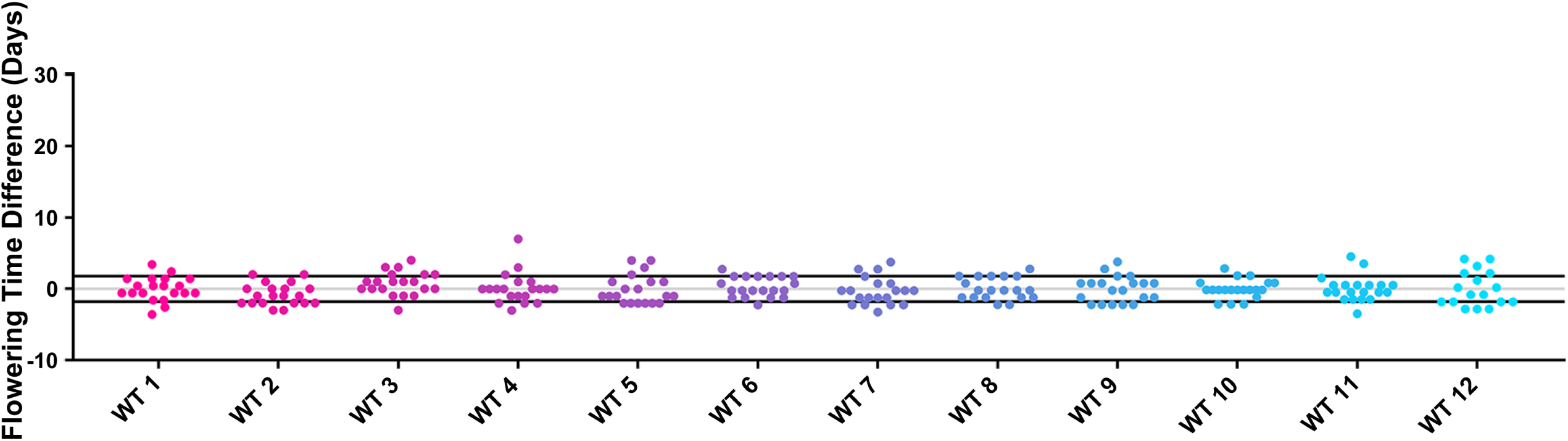
1 cm Bolting Age Distributions of Control Plants. Values presented are the difference between the age at 1 cm inflorescence of the individual control plant and the average age at 1 cm inflorescence of the control in the accompanying experiment. The grey line is at the average control value and the black lines are at +/- the standard deviation of the control plants.

**Figure 3 – Figure Supplement 1.**
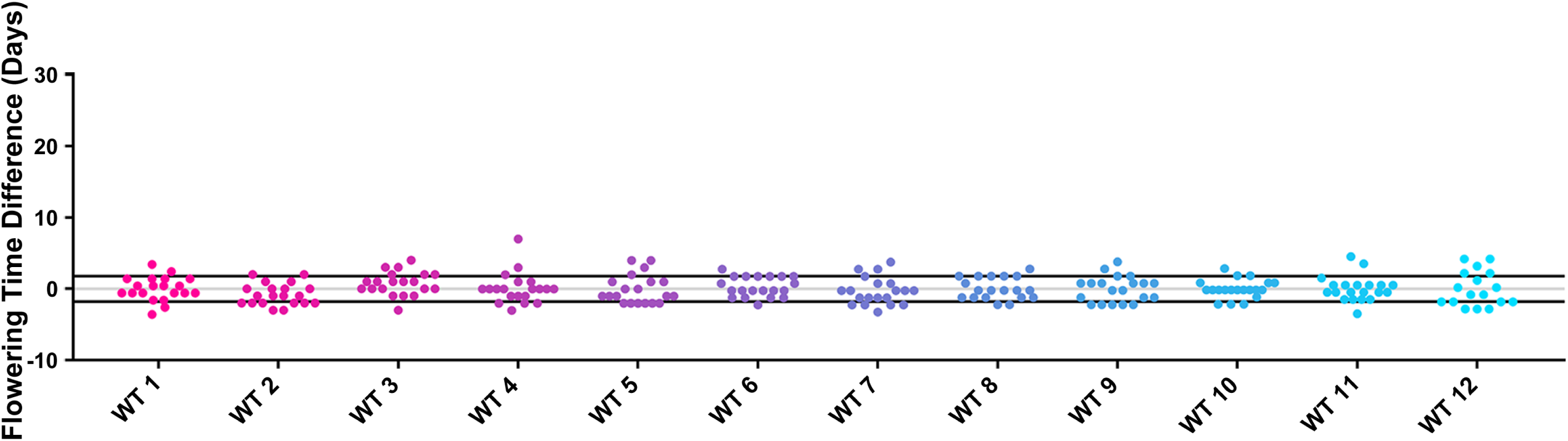
Anthesis Delay Distributions of Control Plants. Values presented are the difference between the anthesis delay of the individual control plant and the average anthesis delay of the control in the accompanying experiment. The grey line is at the average control value and the black lines are at +/- the standard deviation of the control plants.

**Figure 4 – Figure Supplement 1.**
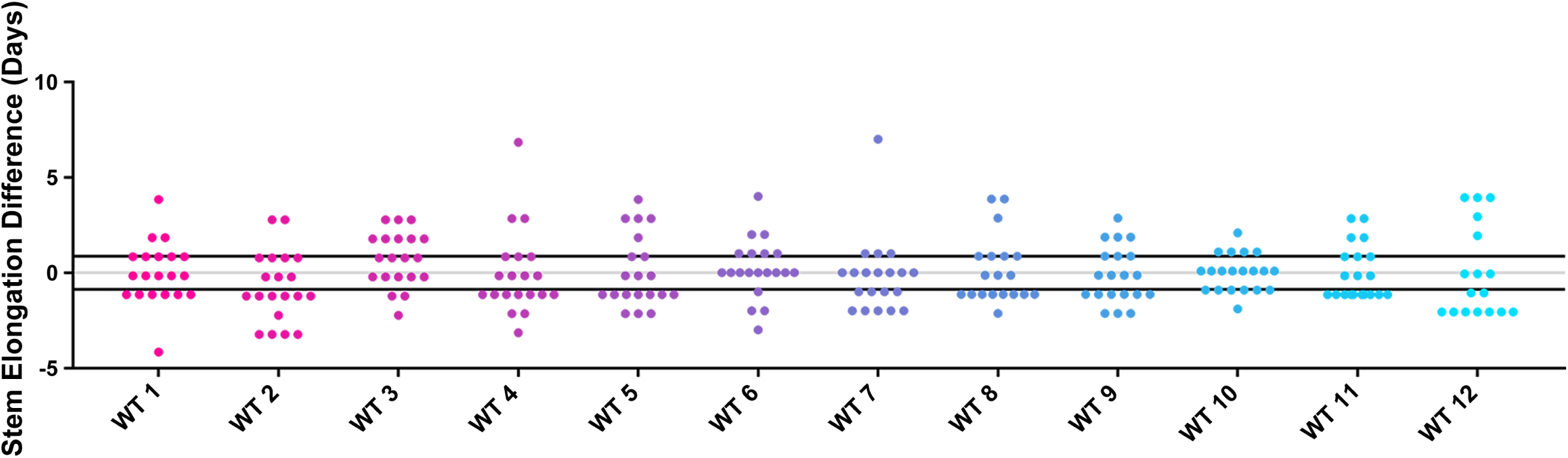
Stem Elongation Time Distributions of Control Plants. Values presented are the difference between the stem elongation time of the individual control plant and the average stem elongation period of the control in the accompanying experiment. The grey line is at the average control value and the black lines are at +/- the standard deviation of the control plants.

**Figure 6 – Figure Supplement 1.**
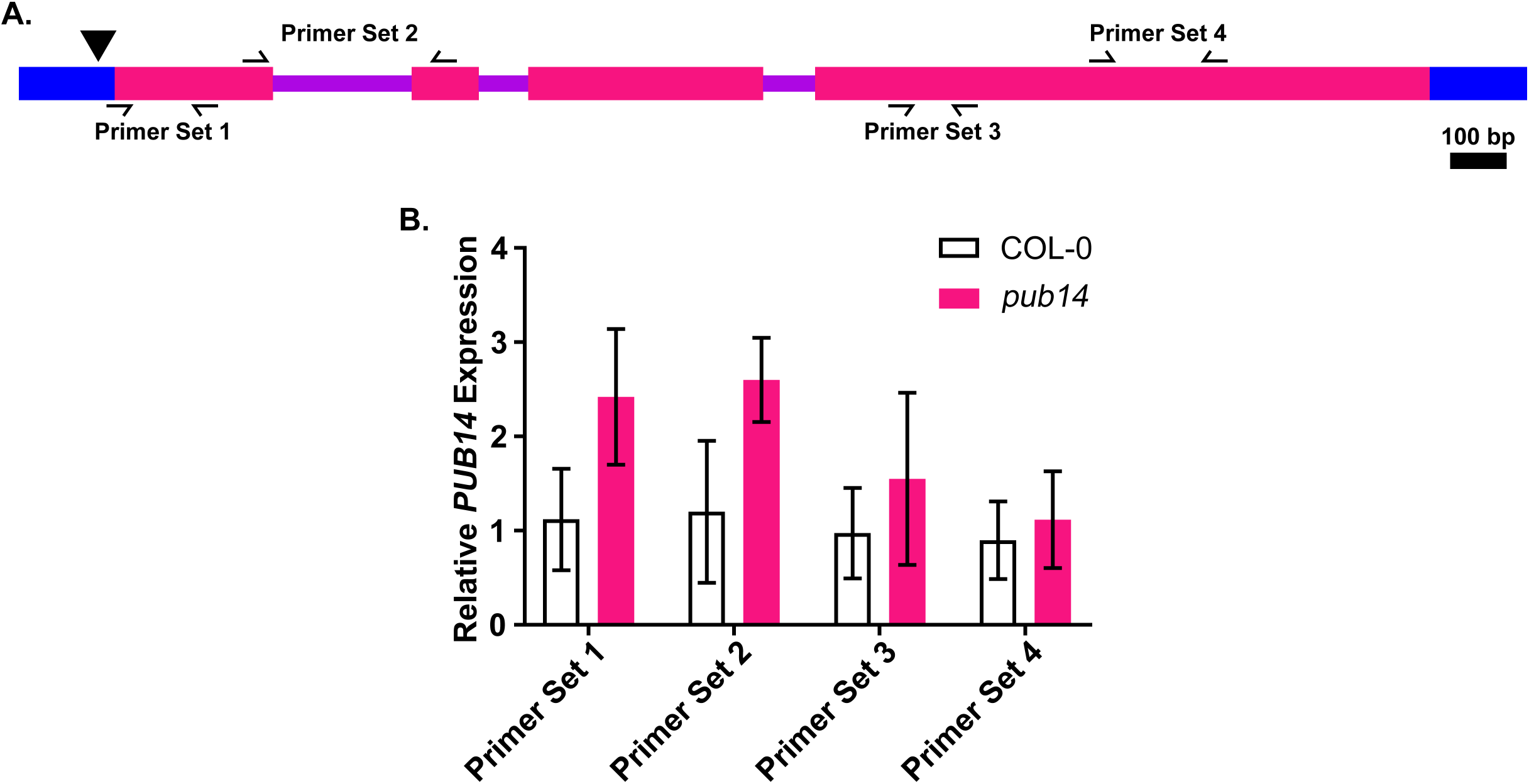
*PUB14* expression in the *pub14-1* mutant. A) Diagram of the genomic structure of *PUB14*. Blue represents 3’ and 5’ UTR sequences, pink represents exon sequences, purple represents intron sequences. The black triangle represents the T-DNA insertion location. Black arrows represent primer locations. B) *PUB14* expression was measured using quantitative RT-PCR in wild type or homozygous *pub14-1* mutants grown under long day (16 hours light/8 hours dark) conditions. Quantifications are the average of three biological replicates with error bars showing standard deviation.

Table S1. IP-MS Results from the PUB14 decoy.

Table S2. Source Data for Figures 1-4 and their Supplements

